# CpG-ODN 2007 protects zebrafish against *Vibrio vulnificus*-induced infection

**DOI:** 10.1101/780742

**Authors:** Hua Chen, Lijuan Zhang, Suyi Li, Ling Ke, Chentao Lin, Xu Chen

## Abstract

*Vibrio vulnificus (V. vulnificus)* is an aquatic pathogen that can cause primary sepsis and soft tissue infection. CpG oligodeoxynucleotides (CpG-ODN) are a type of essential immunomodulators, which can trigger or enhance immune responses in mammals, fish, and humans. In this study, we evaluated the effect of CpG-ODN 2007 as a potential immunostimulant for zebrafish infected with *V. vulnificus* (FJ03-X2). Fish injected with the CpG-ODN 2007 showed lower mortality rate compared with the fish that did not receive treatment. The survival rates of CpG-ODN 2007-treated group and PBS-treated group were 85% and 57.9%, respectively. In addition, our *in vitro* results demonstrated that CpG-ODN 2007 can effectively reduce the toxicity of *V. vulnificus* (FJ03-X2) to zebrafish embryonic fibroblast (ZF4) cells. Furthermore, we assessed immune-related genes expression patterns in FJ03-X2 infected zebrafish or ZF4 cells with and without CpG-ODN 2007 treatment, such as TLRs and IL-1β. To sum up, our data indicated that CpG-ODN 2007 protects zebrafish against *Vibrio vulnificus* induced infection.

## Introduction

Intensive culture systems and overcrowding of cages in a limited area might cause adverse effects on aquaculture (Bunlipatanon and U-taynapun 2017). Besides, disease outbreaks in fish are very frequent and often result in massive economic losses, which restrict the sustainable development of aquaculture industry (Ding et al. 2018). Currently, antibiotics are still used as the first treatment choice to fight bacterial infections. Nevertheless, the antibiotic resistance is accelerated by overuse of antibiotics (Ding et al. 2018). Therefore, it is urgent and necessary to develop alternative, effective and safer means to control bacterial diseases progression and spreading.

*Vibrio vulnificus* is a Gram-negative pathogenic bacterium widely distributed in seawater, plankton, seabed sediments and aquatic products (Hernandez-Cabanyero et al. 2019). It is extremely pathogenic for humans and lower vertebrates, including amphibians, reptiles, since it can lead to primary sepsis, soft tissue infection and extensive cellular damage (Lee et al. 2009). *V. vulnificus* is a prominent invasive bacterial species within the Vibrio genus, and its infection has become an important health problem in aquaculture (Liu et al. 2019). So far, *V. vulnificus* infections have been observed in the eel *Anguilla Anguilla* (AUSTIN et al. 1999), salmon *Salmo salar* (Lunder et al. 2000), sea bass *Dicentrarchus labrax* (Abdullah et al. 2017), trout *Oncorhynchus mykiss* (Pedersen et al. 2008), and Chinese shrimp *Penaeus chinensis* (Teng et al. 2017). In fish, the infection can cause external haemorrhages and ulceration of the skin, and internal haemorrhages in the intestines, nodular lesions in the spleens and kidneys (Liu et al. 2019). In addition, the host response to *V. vulnificus* infection by overproduction of TNF-α, IL-1β, and other inflammatory cytokines can cause endotoxemic shock and lead to high mortality (Pan et al. 2011).

CpG oligodeoxynucleotides (CpG-ODN) are a type of essential immunomodulators, which can trigger or enhance immune responses in mammals, fish, and humans (Jung and Jung 2017). Cytokines are one of the major regulators of immune responses (Tassakka et al. 2006). CpG-ODNs can be recognized as pathogen-associated molecular pattern (PAMP) of microbial DNA by Toll-like receptor 9 (TLR9) on B-cells, dendritic cells (DCs), natural killer (NK) cells, and monocytes; and the resulting cascade signaling can induce pro-inflammatory cytokine production and activate immune protection mechanisms to the host (Nguyen et al. 2017). CpG-ODN can induce protection against different bacterial diseases in various fish species, such as *Aeromonas salmonicida* in rainbow trout, *Streptococcus iniae* in striped bass, amoebic gill diseases in Atlantic salmon, and *Edwardsiella tarta* in olive flounder (Pridgeon et al. 2012). The immune responses induced by CpG ODNs in fish are manifold, including macrophage activation, leucocyte proliferation, and cytokine secretion (Liu et al. 2010).

CpG-ODN 2007 can induce cell proliferation in goats, IFN-g production, and balanced immune responses in cattle (Ioannou et al. 2002). In addition, CpG-ODN 2007 has been shown to elicit strong immune responses against infectious pathogens in fish primarily through innate immune responses (Pridgeon et al. 2012). Cha *et al* have confirmed the immunostimulatory and protective effects of CpG-ODN 2007 against *Edwardsiella tarda* infection in olive flounder (Cha et al. 2017). Furthermore, Julia *et al* found that CpG-ODN 2007 with an adjuvant such as QCDCR can be used to protect *Nile tilapia* against *S. iniae* infections (Pridgeon et al. 2012). In addition, CpG-ODN 2007 administration has shown to exert a significant effect against *Streptococcus iniae* in tilapia (Cha et al. 2017). However, whether CpG-ODN 2007 is capable of constraining the *V. vulnificus* infection in zebrafish has not been yet reported.

In the present study, we examined the effect of CpG-ODN 2007 as a potential immunostimulant for zebrafish infected with *V. vulnificus* (FJ03-X2) and evaluated its exact mechanism of action, thus laying a foundation for vaccine development and disease prevention of important economic fish.

## Results

### CpG-ODN 2007 showed protection effect against *V. vulnificus* infection

After 12 hours of infection with FJ03-X2, some zebrafish began to show typical clinical symptoms, including swollen abdomen, abdominal fin and caudal fin hemorrhages, and eventually died later on (Figure 1A,B).

**Fig. 1.**
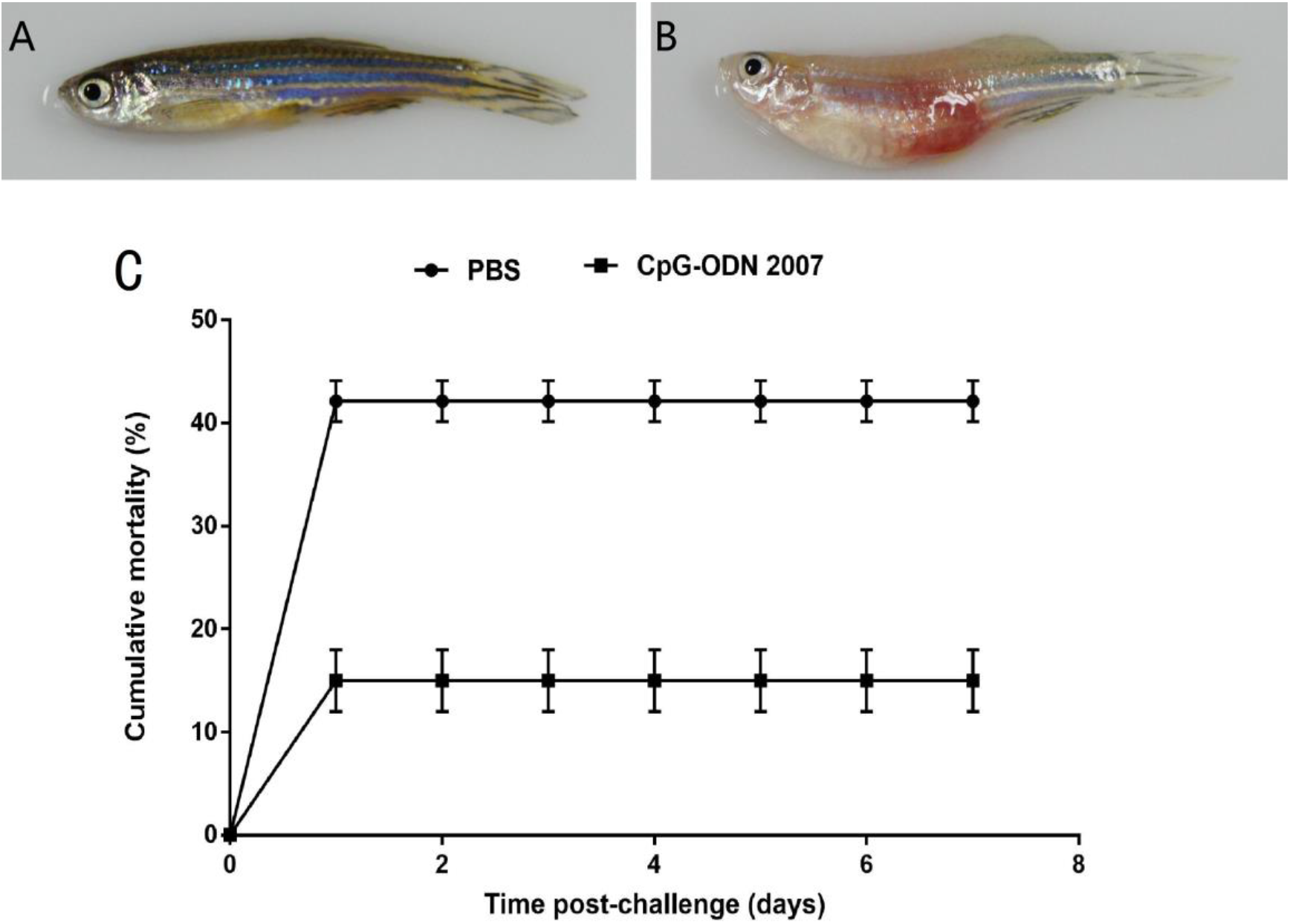
Immune protection test by CpG-ODN 2007. (A) Uninfected healthy zebrafish. (B) Morphological changes of zebrafish infected with V. vulnificus. (C) Cumulative mortality of immunized groupers after V. vulnificus challenge.

Cumulative mortality of zebrafish was recorded from day 1 post infection until the end of the experiment. Briefly, in all groups, no death was observed at day 1; the mortality rate remained constant for up to one week. The cumulative mortality of the control (PBS-treated fish) was 42.1%, whereas that of CpG-treated fish was 15.0%. By calculation, the relative percentage survival (RPS) of the CpG-ODN 2007 group was 64.37%. These data suggested that CpG-ODN 2007 has a protective effect and can significantly improve the anti-invasive Vibrio ability of zebrafish (Figure 1C).

### Expressions of immune-related genes in the intestine

Mucosal-associated immune tissue in the intestine is important immune organ in fish, and the first line of host defense system against infection.

The mRNA levels in the intestine of two groups of zebrafish are shown in Figure 2. The results showed that the expressions of all genes were significantly downregulated in the CpG-treated group compared to the PBS-treated group (P<0.01). Among them, the expression level of IL-1β, which was 99.88% lower than that of the control group, was the most significant.

**Fig. 2.**
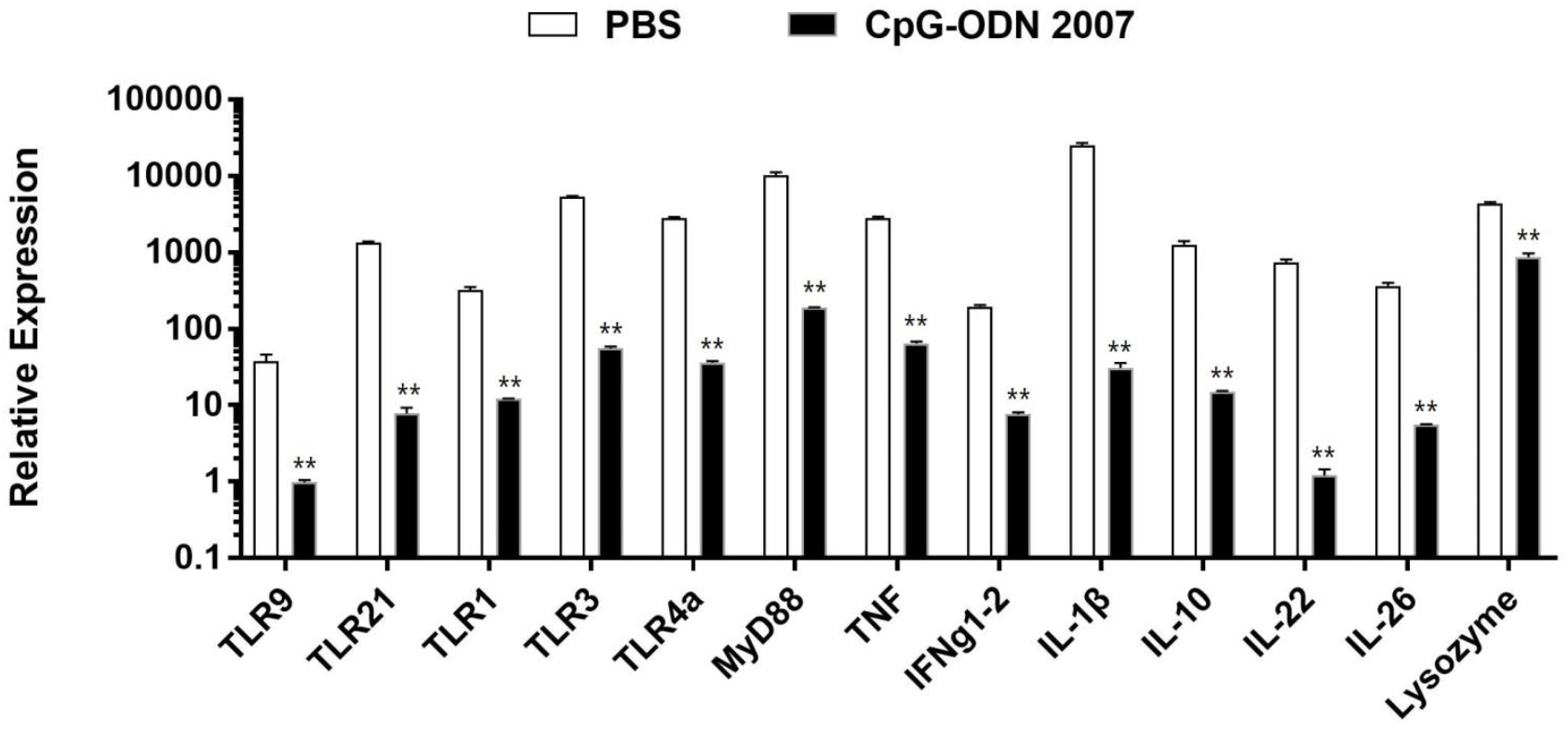
Effect of V. vulnificus (FJ03-X2) infection on the expression levels of immune-related genes in the intestine of zebrafish. Expression levels of TLR9, TLR21, TLR1, TLR3, TLR4a, MyD88, TNF, IFNg1-2, IL-1β, IL-10, IL-26, IL-22 and lysozyme analyzed using Real-time RT-PCR. The relative expression is the mRNA expression value for the target gene β-actin. Data are presented as the mean S.E. **P<0.01.

### Expressions of immune-related genes in the kidney

The kidney is one of the main immune organ of fish; the main site for the growth and differentiation of fish immune cells, and the main part of the immune response.

The mRNA levels in the kidney of two groups of zebrafish are shown in Figure 3. Compared to the control group, TLR9, lysozyme and IL-26 were significantly upregulated, while IL-1β was significantly downgraded (P<0.01) in the kidney of the CpG-treated group (P<0.01). No differences in other genes were observed between groups.

**Fig. 3.**
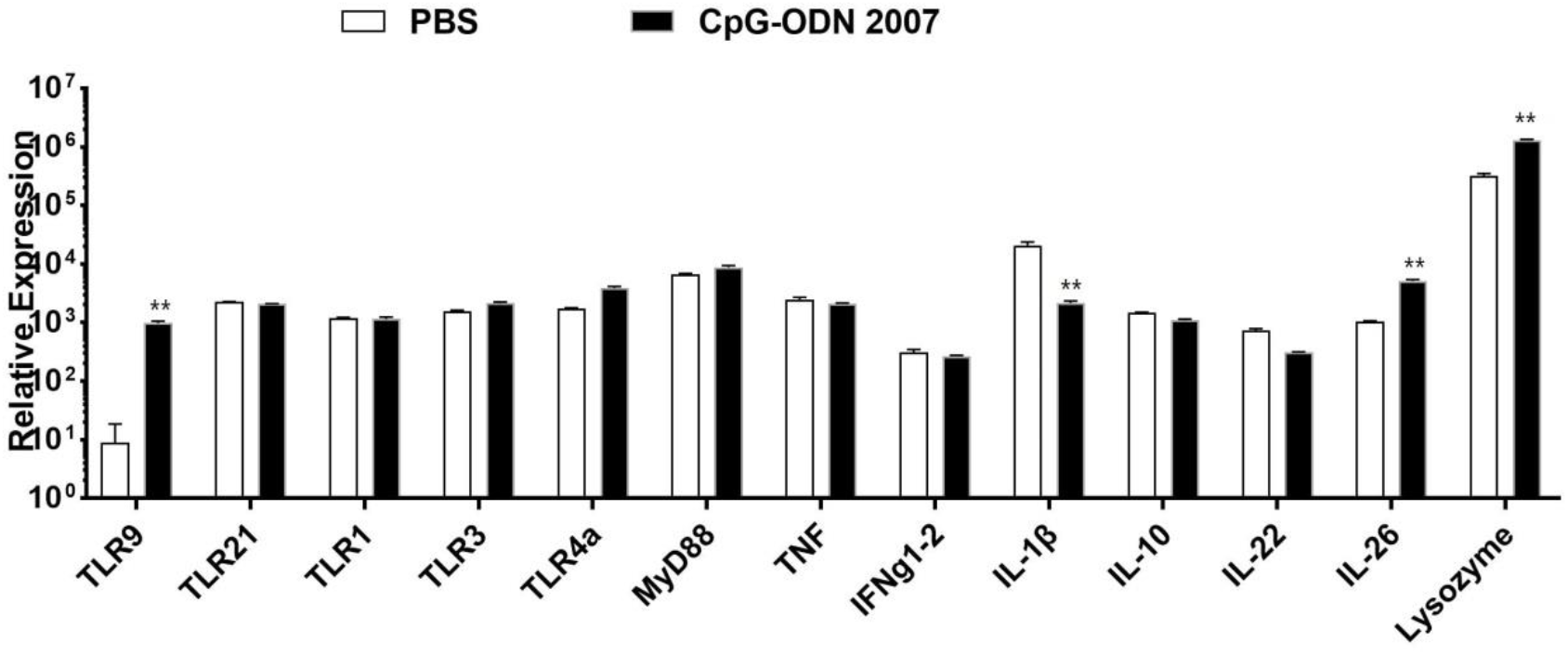
Effect of V. vulnificus (FJ03-X2) infection on the expression levels of immune-related genes in the kidney of zebrafish. The expression levels of TLR9, TLR21, TLR1, TLR3, TLR4a, MyD88, TNF, IFNg1-2, IL-1β, IL-10, IL-26, IL-22 and lysozyme analyzed using Real-time RT-PCR. The relative expression is the mRNA expression value for the target gene β-actin. Data are presented as the mean S.E. **P<0.01.

### Expressions of immune-related genes in the gill

Along with kidneys and intestine, gills are the first line of defense against pathogens.

The mRNA levels in the gill of two groups of zebrafish are shown in Figure 4. The expression level of TLR9 was 3.7-fold higher than that of the PBS group (P<0.01). In addition, the genes of MyD88 and IL-1β were significantly downgraded in CpG-treated group compared to PBS-treated (P<0.01), while no difference in other genes were observed between

**Fig. 4.**
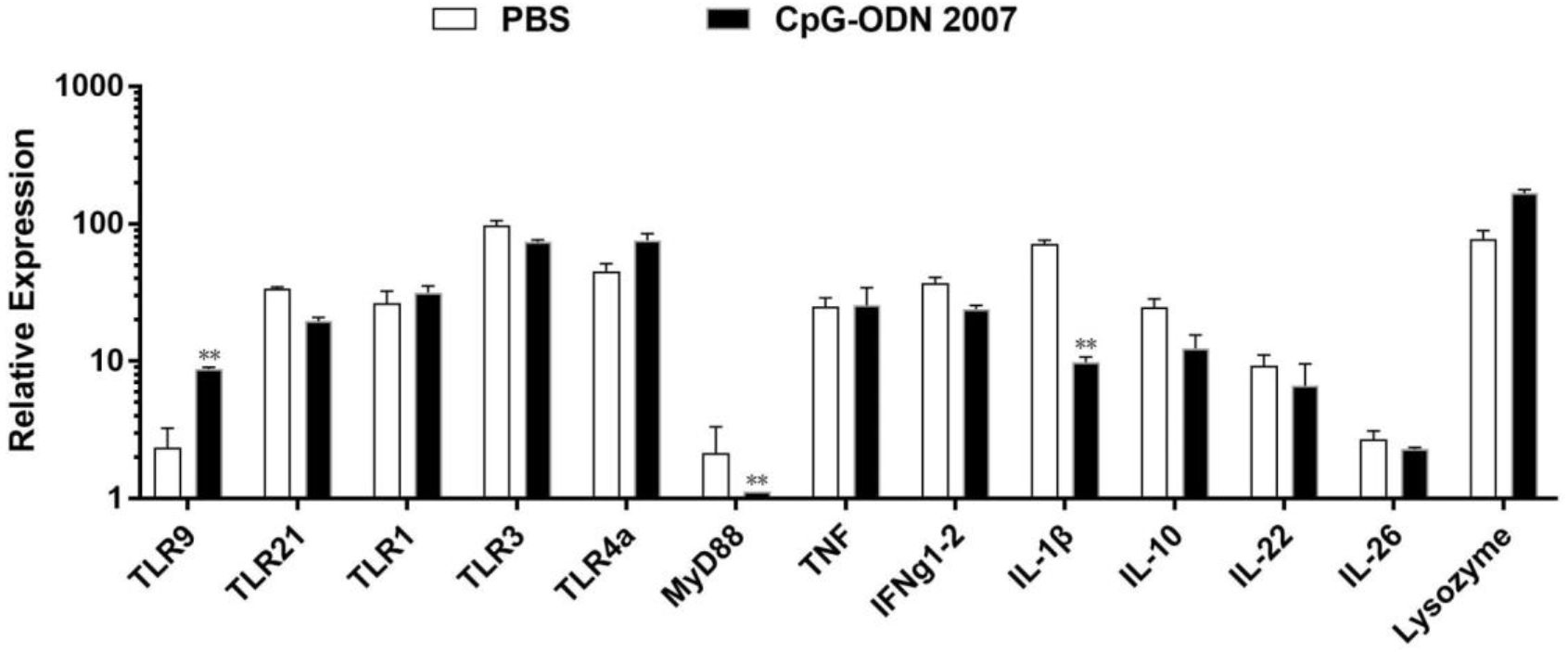
Effect of V. vulnificus (FJ03-X2) infection on the expression levels of immune-related genes in the gills of zebrafish. Expression levels of TLR9, TLR21, TLR1, TLR3, TLR4a, MyD88, TNF, IFNg1-2, IL-1β, IL-10, IL-26, IL-22 and lysozyme analyzed using Real-time RT-PCR. The relative expression is the mRNA expression value for the target gene β-actin. Data are presented as the mean S.E. **P<0.01.

### CpG-ODN 2007 help ZF4 cells resist *V. vulnificus* infection

To further explore the antibacterial mechanism of CpG-ODN 2007, zebrafish embryonic fibroblast cell line (ZF4) was used to evaluate the antibacterial ability of CpG-ODN 2007. We examined the cytotoxicity of FJ03-X2 in ZF4 cells with or without CpG-ODN 2007 treatment. As shown in Figure 5, we found similar lactate dehydrogenase (LDH) level in ZF4 cells treated with CpG-ODN 2007 and those that did not receive CpG-ODN 2007 treatment (10.81% and 6.71%, respectively) which further suggested that CpG-ODN 2007 is non-toxic to ZF4 cells. However, compared with uninfected ZF4 cells, the cytotoxicity of the two groups after the infection was significantly increased, indicating that the FJ03-X2 strain is highly toxic (CpG-untreated group: 88.10%; CpG-treated group: 54.61%; contour groups: <10%; however, the cytotoxicity was significantly lower in the CpG-treated group compared to CpG-untreated group (all P<0.01). These results suggested that pretreated CpG-ODN 2007 pretreated might help the ZF4 cells resist FJ03-X2 infection.

**Fig. 5.**
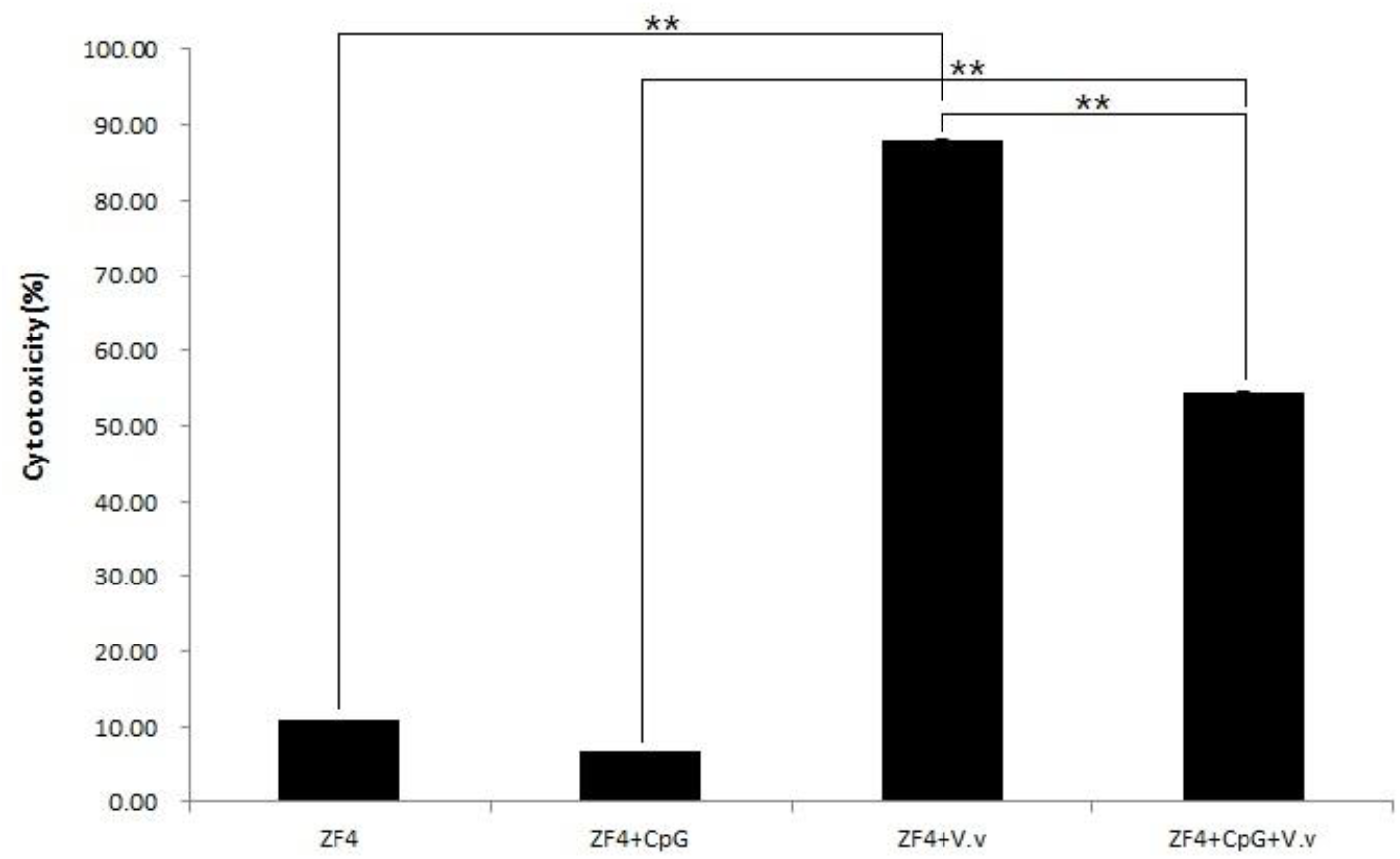
Cell Cytotoxicity in non-infected and infected ZF4 cells. **P<0.01.

### Immune-related genes expression in *V. vulnificus* infected ZF4 cells

The mRNA levels after FJ03-X2 strains infection are shown in Figure 6. Compared with CK and EK, we found that the expression of immune-related genes in ZF4 cells was significantly increased after stimulation with CpG-ODN 2007, which indicated that CpG-ODN 2007 could induce the signal response of these genes and activate the immune response in ZF4 cells.

**Fig. 6.**
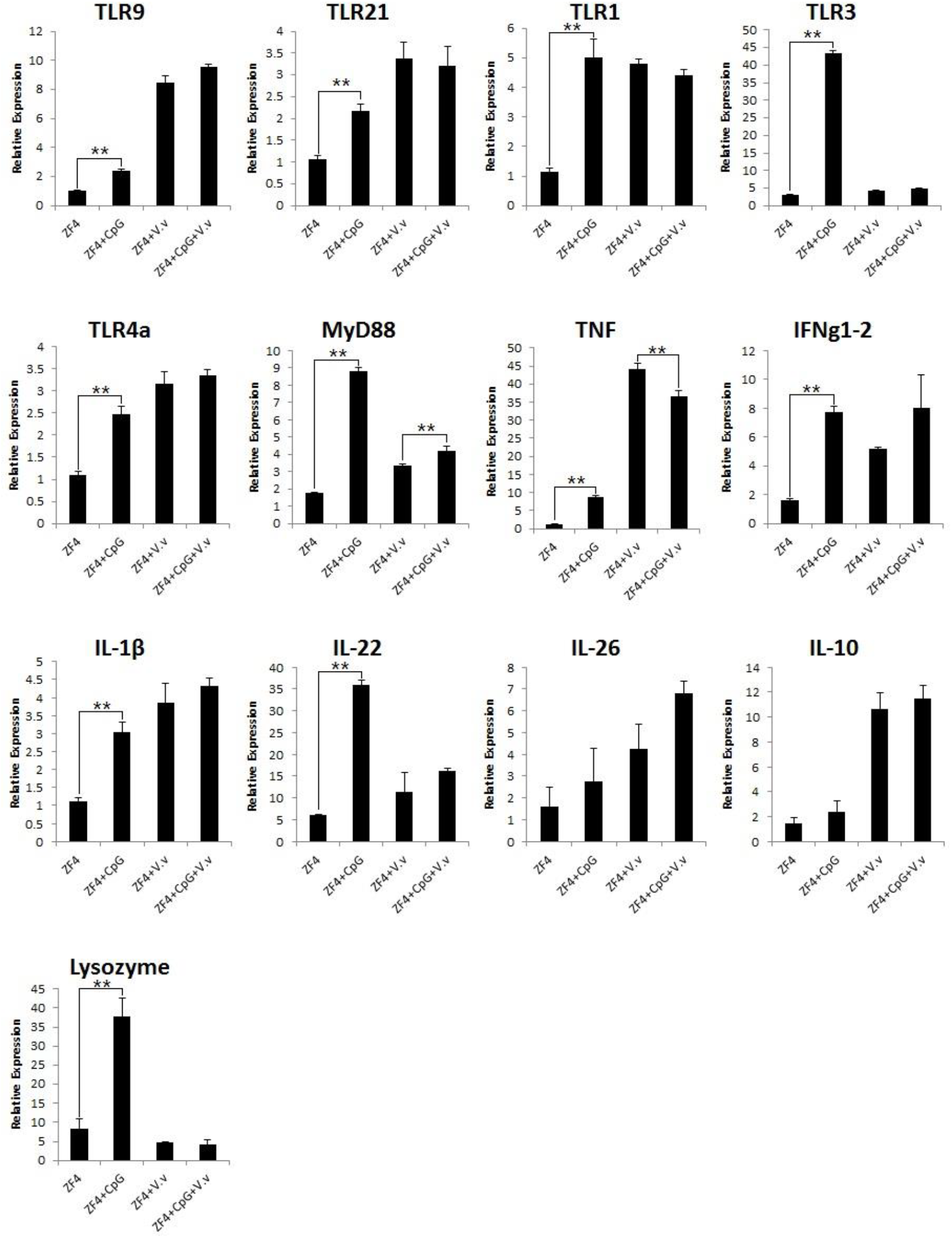
Expression levels of immune-related genes in non-treated and CpG-ODN 2007-treated groups of zebrafish before and after V. vulnificus infection. CK (ZF4), EK (ZF4+CpG), CV (ZF4+V.v) and EV (ZF4+CpG+V.v). The relative expression is the mRNA expression value for the target gene β-actin. Data are presented as the mean □ S.E. **P<0.01.

Compared with CV and EV, ZF4 cells were not significantly different in the control group and the experimental group after FJ03-X2 strains infection (except MyD88, TNF). However, by comparing CK and CV, the gene expression of ZF4 cells was significantly increased after FJ03-X2 strains infection (except TLR3, Lysozyme), which indicated a strong inflammatory response. Compared with EK and EV (CpG-ODN 2007-treated group), the expression levels of TLR9, TLR21, TLR4a, TNF, IFNg1-2 and IL-1β were upregulated after FJ03-X2 infection, but the upregulation was much smaller than that of the non-treated group; the expression levels of TLR1, TLR3, MyD88 and IL-22 genes were down-regulated. This indicated that CpG-ODN 2007 can inhibit overexpression of these genes when infected with *V. vulnificus*.

## Discussion

In this study, we evaluated the effect of CpG-ODN 2007 as a potential immunostimulant for zebrafish infected with *V. vulnificus* (FJ03-X2). We found that CpG-ODN 2007 is an effective adjuvant, which could be used to fight *V. vulnificus* infection in zebrafish. In addition, we discovered that CpG-ODN 2007 could effectively reduce the toxicity of *V. vulnificus* to zebrafish embryonic fibroblast (ZF4) cells. These studies further our understanding of the innate immune system of fish, which can be used as a reference for the application of CpG-ODN in aquaculture.

In our previous study, we examined the expression of related immune factors (TLR9, TLR21, TLR1, TLR3, TLR4a, MyD88, TNF, IFNg1-2, IL-1β, IL-10, IL-26, IL-22 and Lysozyme) before and after injection of CpG in zebrafish, and found that the expression levels of these genes were upregulated in CpG-ODN 2007 treated-group (data not shown). Similarly, Jung *et al* discovered that gene expression of TLR9, MyD88 and IL-1β were significantly elevated in the CpG-ODN1668 treated group (Jung and Jung 2017). Moreover, Nguyen and his team found that CpG-ODN1668 displays higher activation capacity of TLR9 and TNF as compared to the controls (Nguyen et al. 2017); CpG-ODNs stimulated the expression of immune genes in the head kidney of common carp (Tassakka et al. 2006). In this study, we evaluated the immunostimulant effects of CpG-ODN 2007 in zebrafish *in viv*o. Fish injected with CpG-ODN 2007 showed higher survival rate compared to untreated fish challenged with *V. vulnificus*, which suggests that CpG-ODN 2007 has a protective effect and can effectively improve the antibacterial ability of zebrafish. Next, we analyzed the expression patterns in the intestine, kidney and gills following *V. vulnificus* infection, which are the first line of defense against pathogens (王俊丽 et al. 2014). We found that the expression of immune-related genes was most significant in the CpG-treated and non-treated group after FJ03-X2 infection. We also speculated that the intestine might be the main organ of the acute infection period of *V. vulnificus* in zebrafish.

*V. vulnificus* is an extremely virulent bacterium that contains a virulence factor responsible for the regulation of intestinal colonization, which often causes acute inflammatory responses and the killing of phagocytes in the intestine (Lee et al. 2017). In the intestine, the expression levels of all tested genes in the CpG-treated group were significantly lower than those in the PBS-treated group after *V. vulnificus* infected. Among them, IL-1β was the most significant and its expression level was only 0.12% of the control group. The virulence factor (VvpM) generally targets the critical pathogenic pathway of *V. vulnificus* to promote IL-1β production coupled with necrotic macrophage death (Lee et al. 2017). Moreover, CpG-ODN 2007 may regulate the over-expression of IL-1β to avoid damage to the host. The Toll-like receptors (TLRs) are pattern recognition receptors (PRRs) that are strongly associated with both the innate immune and adaptive immune systems (Bilodeau-Bourgeois et al. 2008). As the powerful first line of defense in response to pathogen invasion, TLRs can recognize specific components of microorganisms and transmit signals to NF-κB or MAPK via MyD88, thereby producing cytokines such as TNF-α, IL-1β and IFN involved in immune regulation (Nguyen et al. 2017). In this study, the expression of TLRs in the intestine was significantly lower than that in the control group. The upregulation of TLR expression indicates the process of immune clearance of pathogens when the body is infected by bacteria (费超 2013). Once the residual bacteria are removed, the TLR level decreases. Considering that the expression of TLRs in the intestine was significantly lower in the CpG-treated group, we speculated that the bacteria in the intestine were cleared, and expression status was turned off, which however was not the case in the control group. These data suggest that CpG-ODN 2007 can promote the host to produce an immune response and eliminate pathogenic bacteria.

MyD88, as an adaptor, is significantly downregulated in the intestine that is affected by the expression level of TLRs. In this study, we found that the downregulation of TLRs reduced the activation of TLR-MyD88 signaling pathway, accompanied by decreased secretion of downstream molecules such as IFNg1-2 and IL-10. TLR signaling pathway can induce the expression of a variety of pro-inflammatory factors, thereby triggering the inflammatory response of the fish, and producing a warning signal for the invasion of pathogens (Li et al. 2016). Similar to IL-1β, TNF-α is a potent pro-inflammatory cytokine in early inflammatory events (Muire et al. 2017). Controlled TNF-α has critical immunoregulatory roles while its overproduction is injurious as it results in inflammatory diseases (Wang et al. 2011). In this study, the expression of TNF was significantly downregulated in CpG-treated group that survived after infection. These data suggest that CpG-ODN 2007 protects the body by regulating the overexpression of TNF.

In the kidneys and gills, the expression of TLR9 was significantly upregulated and IL-1β was significantly downregulated, which is consistent with previous studies (Jorgensen et al. 2001). It is speculated that CpG-ODN 2007 is mainly recognized in these two organs where it may reduce the overexpression of IL-1β (Tassakka et al. 2006). Lysozyme is considered as one of the key components for the innate immune response to pathogen infection with their strong antibacterial properties, and its expression level is considered an important index for monitoring fish immunity (Gao et al. 2016). Interestingly, in this study, the expression of lysozyme was significantly upregulated in CpG-treated group that survived after infection in the kidney and gill, while shows the opposite expression pattern in the intestine. This suggested that lysozyme has different expression patterns in different tissues, which is consistent with the study results reported by Gao et al (Gao et al. 2016).

Furthermore, our *in vitro* experiments showed that CpG-ODN 2007 can effectively reduce the toxicity of *V. vulnificus* to ZF4 cells. After CpG-ODN 2007 stimulation, the expression of immune-related genes in ZF4 cells was significantly increased, indicating that CpG-ODN 2007 could induce the signal response of these genes and activate the immune response, which is consistent with Tassakka *et al* (Tassakka et al. 2006). We also found that the expression of TLR21, TLR1, TNF, IL-1β and other inflammation-related factors was suppressed after treatment with CpG-ODN 2007, which was not observed in the control group. These results suggest that CpG-ODN 2007 can regulate the immune system to control overexpression of certain genes. We also speculate that this might be the cause of the reduced cytotoxicity, which needs to be confirmed by future studies.

In conclusion, this is the first study that investigated the ability of CpG-ODN 2007 to increase resistance against *V. vulnificus* (FJ03-X2) challenge in zebrafish. The study expands the current knowledge of the effect of CpG-ODN 2007 on the immune system of fish by examining the expression levels of immune-related gene. *In vivo* experiments showed that CpG-ODN 2007 stimulation can effectively improve the survival rate of zebrafish infected with *V. vulnificus*. Furthermore, the *in vitro* experiments showed that CpG-ODN 2007 can reduce the toxicity of *V. vulnificus* to ZF4 cells. Our study confirmed that CpG-ODN 2007 can significantly enhance immune responses, and provide effective protection against *V. vulnificus* in zebrafish, thus having a great potential as an immunostimulant in aquaculture. Certainly, more work is required to optimize the usage of CpG ODN in fish.

## Materials and Methods

### CpG-ODN 2007

CpG-ODN 2007 (5’-TCGTCGTTTTGTCGTTTTGTCGTT-3’) were synthesized by Takara Biotechnology (Dalian) Co. Ltd..

### Bacterial strains

*V. vulnificus* strain FJ03-X2 was isolated from European eel by Fish Disease Research Laboratory of Institute of Biotechnology, Fujian Academy of Agricultural Sciences in 2003(许 斌 福 et al. 2005). The strain possess strong virulence, good high immunogenicity and wide fish applicability (许斌福 et al. 2010).

### Cell lines

ZF4 cell were purchased from China Zebrafish Resource Center. Cells were cultured in DME/F12 medium (Hyclone, No.SH30023.01, USA) supplemented with 10% fetal bovine serum (GIBCO BRL, Grand Island, NY, USA) and antibiotics (10 unit⋅mL^−1^ penicillin G and 10 μg⋅mL^−1^ streptomycin; GIBCO, No.15140, USA) in a humidified atmosphere containing 5%CO_2_/95% air at 28ºC.

### Zebrafish

Wild-type strains of zebrafish (AB) (>6 months, weighing 0.2g on average) were purchased from China Zebrafish Resource Center. All the animals were cultivated in a re-circulated water aquatic animal culturing system (self-made) at 28 °C. The circulating water was constantly filtered, and the salinity was 0.05%. Fish were fed with 1% body weight twice daily using commercial dry fish food. All animal studies were done in compliance with the regulations and guidelines of Fujian Academy of Agricultural Sciences animal care and conducted according to the AAALAC and the IACUC guidelines.

### Zebrafish immunization and *V. vulnificus* infection

Zebrafish were acclimated for one week before experiments and fasted one day before treatment. Adult zebrafish were then injected with 10 μL of CpG-ODN 2007 (1μg⋅tail^−1^) or PBS, respectively, and cultured in two tanks (n=20 fish per pool, three repetitions per group). Two days post injection, zebrafish were challenged with LD_50_ of *V. vulnificus* by intraperitoneal injection at 7.36× 10^3^ cfu per fish (陈华 et al. 2018). Cumulative mortality of the challenged fish was recorded daily for one week. The survival ratio of each treatment was calculated (RPS) using the following formula: *RPS = [1 – (mortality of test group fish%/mortality of PBS control fish%)] × 100*. Intestine, kidneys and gills of randomly selected survival fish were collected and stored at −80℃ for gene expression analysis.

### ZF4 cells infected with FJ03-X2

ZF4 cells were dispensed in 6-well culture plates (2.0×10^5^ cells⋅well^−1^) and then cultured for 24 hours in antibiotic-free growth medium. Cells were then divided into four groups: ①CK, non-treated cells without *V. vulnificus* infection (ZF4); ②EK, CpG-treated cells without *V. vulnificus* infection (ZF4+CpG); ③ CV, non-treated cells with *V. vulnificus* infection (ZF4+*V.v*); ④EV, CpG-treated cells with *V. vulnificus* infection (ZF4+CpG+*V.v*). EK and EV were treated with CpG-ODN 2007, final concentration of 0.01μg⋅μL^−1^ (陈 华 et al. 2018). These cells were continuously cultured for another 48 hours, then used for bacterial infection.

FJ03-X2 were grown overnight at 28°C in TSB medium, centrifuged for 5 min at 8,000 rpm, resuspended and adjusted to 1×10^6^ cfu⋅mL^−1^ in antibiotic-free DMEM-F12 medium. According to previous studies, FJ03-X2 is extremely toxic and can kill cells over a short period of time. When the multiplicity of infection (MOI) is 100:1, the cell death rate can be slowed down, which is useful for observation and detection. In this study, a MOI of 100:1 was selected for testing. The bacterial suspensions were then added to CV and EV. At 24 h post-infection, the supernatants of the cultured cell were collected for cytotoxicity test, while cells were collected and stored at −80℃ for gene expression analysis.

### Cytotoxicity test of *V. vulnificus* to ZF4 cells

The cytotoxicity of *V. vulnificus* was measured by a CytoTox 96® Non-Radioactive Cytotoxicity Assay (Promega Corporation, Madison, WI, USA) according to the manufacturer’s protocol. The lactate dehydrogenase (LDH) released in the supernatants was detected as a marker of cytotoxicity. Briefly, a total of 50μL supernatant from each well of the assay plate was transferred to flatl⍰ bottom 96l⍰ well plate that was prel⍰ loaded with 50 μL⋅well^−1^ reconstituted substrate mix. Following incubation at room temperature for 30 min, 50μL stop solution was added to each well, following by reading the absorbance at 490 nm (Microplate Spectrophotometer, Bio-rad xMark, USA). The cytotoxicity of the *V. vulnificus* was calculated as follows: [(Experimentall⍰ Effector Spontaneous)/ (Target Maximum l⍰ Effector Spontaneous)] x100.

### Real-time PCR (RT-PCR) analysis of gene expression

Total RNA was extracted using TRIzol reagent (Invitrogen, US). RNA concentration was determined via Nano Drop Spectrophotometer ND-1000. cDNA was synthesized using PrimeScript™ RT reagent Kit with gDNA Eraser (TaKaRa, Japan) following the manufacturer’s instructions.

A real-time PCR assay (RT-PCR) was performed using SYBR® Premix Ex Taq™ II Kit to analyze gene expression according to the manufacturer’s instructions, and β-actin was used as the reference gene. The sequences of primers are listed in Table 1. The RT-PCR was performed according to the following conditions: 95℃ for 30s, followed by 40 cycles of 95℃ for 5s, and 60℃ for 34s, followed by a melting curve analysis. Each sample was detected in triplicates. The relative transcriptional levels of measured genes were determined by the 2^−△△Ct^ method.

**Table 1.**
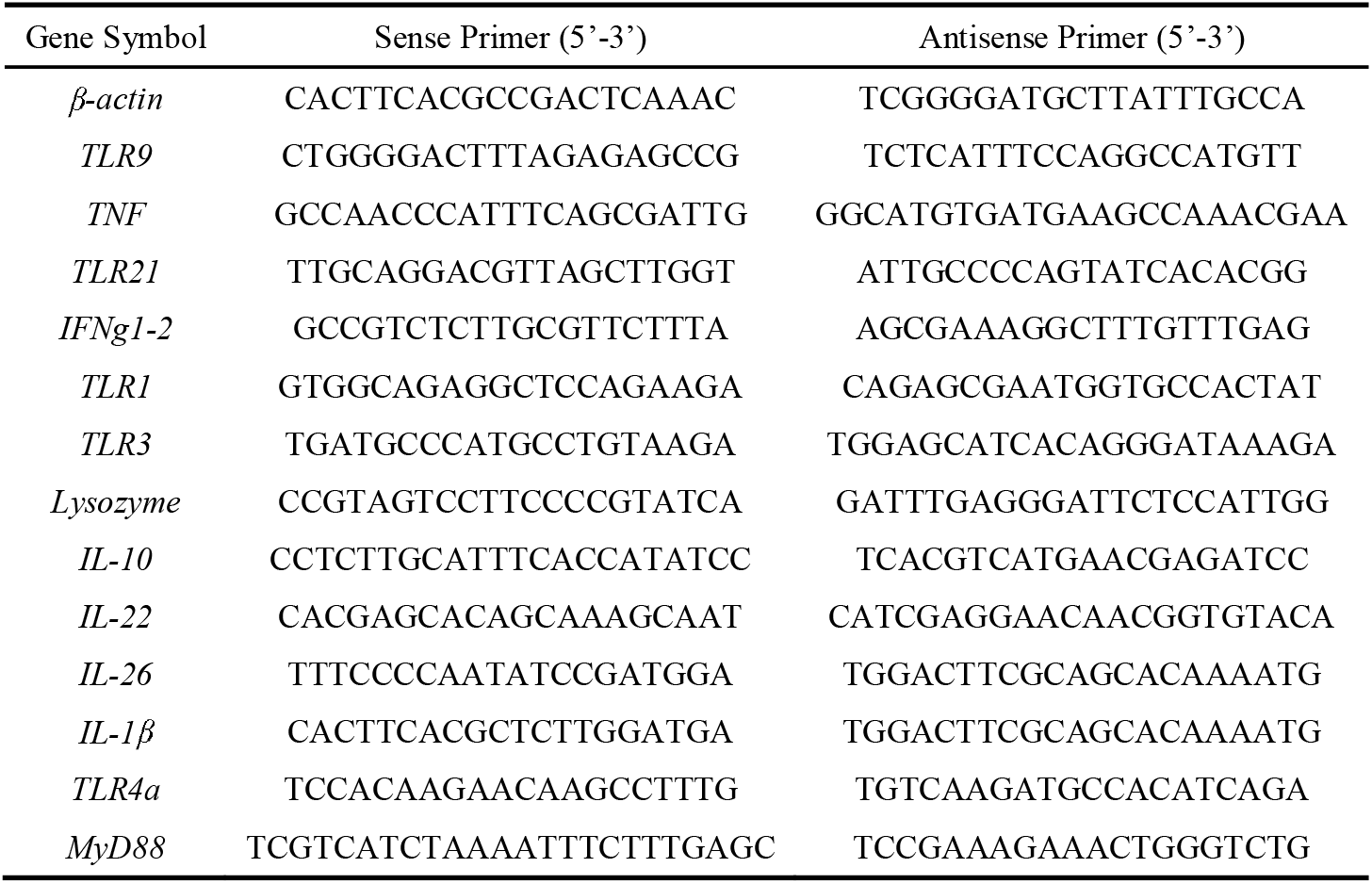
Primers used in this study.

### Statistical analysis

Each experiment was repeated at least three times. Analysis of variance (ANOVA) was used to analyze the differences. Data were presented as means ± standard deviations. A P < 0.01 was considered to be statistically significant.

## Acknowledgements

Not applicable.

## Availability of data and materials

The data set supporting the results of this article are included in the article.

